# Palmitic acid reduces viability and increases production of reactive oxygen species and respiration in rat tendon-derived cells

**DOI:** 10.1101/2023.02.08.527761

**Authors:** Subhajit Konar, Christopher P Hedges, Karen E Callon, Scott Bolam, Sophia Leung, Jillian Cornish, Dorit Naot, David S Musson

## Abstract

Clinically, there is a positive correlation between BMI and the risk of tendinopathy. However, the underlying mechanisms are not understood. Dyslipidaemia and increased circulating free fatty acids (FFA) are associated with increased BMI. We hypothesised that increased FFA concentrations negatively affect rat tendon-derived cells (rTDCs) through mitochondrial-mediated mechanisms.

rTDCs were isolated and treated with oleic acid (OA), stearic acid (SA), and palmitic acid (PA). Cell viability was assessed using AlamarBlue™ assay, and gene expression using real-time PCR. Cell respiration and reactive oxygen species (ROS) production were measured using high-resolution respirometry and MitoSox staining. PA transport into the mitochondria was blocked by pre-treatment with 50µM etomoxir.

Treatment with SA and PA at 10 µg/ml decreased rTDC viability by 40% and 60%, respectively. PA decreased the gene expression of the tendon markers *Scx* and *Tnmd*, and increased the expression of *Mmp3, Mmp13*, and *Ptgs2* (encoding Cox-2). FFA treatment increased the expression of *Cpt1* and *Pdk4*, indicating an increase in mitochondrial FFA oxidation. PA, at 10 µg/ml, increased cellular respiration and ROS production. Pre-treatment with etomoxir partially inhibited the effects of PA on cell viability, *Mmp3* gene expression, ROS production, and cell respiration, but did not affect PA-induced inhibition of *Scx* or *Tnmd* expression.

We found that increased saturated FFA concentrations in the microenvironment reduce cell viability and alter ROS production, respiration, and gene expression. Blocking PA transport into mitochondria partially reversed the negative effects of PA. Overall, an increase in saturated FFA concentrations may contribute to poor tendon health.

## 1 Introduction

Tendinopathy is one of the most commonly diagnosed musculoskeletal disorders and is associated with pain and loss of mobility (Andarawis-Puri et al., 2015). The underlying causes of tendinopathy are multifactorial and are still not entirely known. Numerous epidemiological studies have identified obesity as a risk factor for tendinopathy in both upper and lower limb tendons (Abate et al., 2012; Rechardt et al., 2010; Taş et al., 2017; Titchener et al., 2013). Compared to people with BMI within the normal range, people who are overweight (25<BMI<30) have a 2.6-times greater risk of developing Achilles tendinopathy, and the risk is 6-times greater in people with obesity (BMI>30). Increased BMI is also associated with a thicker tendon and a greater risk of developing plantar fasciitis (Abate et al., 2012).

Increased concentrations of circulating free fatty acids, cholesterol, triglycerides, and glucose in the blood are all hallmarks of obesity (Bi et al., 2019; Boden, 2008; Reaven, 1995). In vitro studies modelling hyperglycaemia and hypercholesterolemia found that these conditions adversely affect tendon cells, and induce a loss of cellular proliferation potential, increased production of reactive oxygen species (ROS), and apoptosis (Li et al., 2020; Lin et al., 2017; Poulsen et al., 2014; Wu et al., 2018). However, the effect of increased concentrations of fatty acids on tendon cells has not been studied. In other tissues, increased saturated FFA concentrations lead to cell death, decreased proliferation, and de-differentiation (Listenberger et al., 2003, Cornish et al., 2008; Elbaz et al., 2010; Ahn et al., 2013; Nemecz et al., 2019). Increased concentrations of FA within cells augment FA mitochondrial oxidation, a process that generates high levels of ROS, which have multiple harmful effects and are associated with cell death (Rosca et al., 2012; Snezhkina et al., 2020; Su et al., 2019).

In this study, we investigated the effects of three of the most abundant circulating FFA: the saturated FFA stearic acid (SA) and palmitic acid (PA), and the monounsaturated oleic acid (OA) (Abdelmagid et al., 2015), on rat tendon-derived cells (rTDCs). rTDC viability and gene expression were assessed in response to different FFA treatments, and etomoxir, which blocks FFA transport into the mitochondria, was used to determine the contribution of mitochondrial processing of FFA to the rTDC effects.

## 2 Material & methods

### 2.1 Isolation of rTDC cells

Primary rTDCs were isolated from Sparge-Dawley rat tails using a previously established protocol (Chhana et al., 2014). Tendon fascicles were pulled from the distal end of the tail, cut into small pieces (<10 mm) and were digested in the digestion mixture overnight at 37°C in a shaker incubator (DMEM/F12, 10% fetal bovine serum (FBS), dispase (0.5 mg/ml) (all from Gibco™, ThermoFisher Scientific Inc.), and collagenase II from *Clostridium histolyticum* (400 units/ml; Sigma Aldrich). Following digestion of the tendon fascicles, the suspension was strained using a 70µm cell strainer (Falcon®, Corning Life Sciences) and resuspended into fresh DMEM/F12-10% FBS media. The cells were cultured in DMEM/F12-10% FBS media in a T-75 tissue culture flask in a bio-incubator at 37°C and 5% CO_2_. Once confluent, the cells were harvested and stored in liquid nitrogen for future use. Before use, passage 1 rTDC were revived and cultured until confluent in a T-75 tissue culture flask and seeded onto 24-well tissue culture plates.

### 2.2 Treatment of tenocyte with fatty acids

rTDCs seeded onto tissue culture plates were incubated overnight in DMEM/F12-5% FBS media at 37°C and 5% CO_2_, and in DMEM/F12-1% FBS for further 24 h. Cells were then treated with 0, 0.1, 1, 10 µg/ml of stearic acid (SA), palmitic acid (PA), and oleic acid (OA) for 24h or 48 h. In experiments that investigated the effect of blocking PA transport into the mitochondria, rTDCs were treated for 2h with 50µM of etomoxir (ETO) before treatment with 10µg/ml PA for 48h (PA+ETO).

### 2.3 Cell viability assay

Cell viability was quantified as previously described (Musson et al., 2015) using alamarblue™ (Invitrogen™, ThermoFisher Scientific Inc.) as a measure of cell number. rTDCs were seeded onto 24-well plates (Greiner BioOne, Sigma Aldrich) at a density of 25,000cells/well, following the above protocol. The effect of fatty acid on cell viability was quantified at 24h and 48h time points. Cell viability was quantified only at 48h for cells treated with PA or PA+ETO. To quantify the cell viability, 5% alamarblue™ (v/v) (Invitrogen, Carlsbad, CA) was added to the culture media and incubated for 4h at 37°C and 5% CO_2_. After the incubation period 200µl of alamarblue™ conditioned media from each well is transformed into 96 well pate (Greiner Bio-One, Frickenhausen, Germany) and fluorescence read (Excitation 540nm, Emission 630nm) in Synergy 2 multi-detection microplate reader (BioTek Instruments, Inc., Winooski, VT). Background fluorescence was subtracted, and the cell viability was normalised to the respective time point control group.

### 2.4 Gene expression RT-PCR

rTDCs were harvested after treatment, and RNA was extracted from the cell pellets by RNeasy® Mini kit (QIAGEN) following the manufacturer protocol, and on-column DNase digestion with the RNAse-Free DNase Set (QIAGEN). Purity and concentration of the extracted RNA were measured using Nano-Drop lite Spectrometer (Thermo-Fisher, Victoria, Australia), with 260/280 absorbance value >1.8 considered acceptable. cDNA was synthesised with super-script-III (Life Technologies, Carlsbad, CA). Gene expression levels were analysed in QuantStudio™ 5 Real-Time PCR System, using multiplex PCR with FAM-labelled TaqMan™ assays for the target genes and VIC-labelled TaqMan™ assay for 18S rRNA, used as the endogenous control (Thermo Fisher Scientific). The relative expression level of the genes compared to control cells (untreated cells at 24hr or relative control of the appropriate experiment as mentioned) was calculated by the 2^-^∆∆Ct method.

### 2.5 Immunofluorescent staining

After 48h of treatment with FFAs, rTDC were stained with Calcein stain (Invitrogen™, Thermo Fisher Scientific) following the manufacturer protocol. Briefly, cells were washed with PBS after 48h of FFA treatment, incubated at 37ºC in 1µM of calcein for 15 mins and visualised under the microscope. Images of the cells were captured using Olympus CKX53 inverted fluorescence microscope using Olympus DP-74 camera.

ROS production by rTDC was quantified by staining with MitoSox^™^ Red (Invitrogen™, Thermo Fisher Scientific). After 48h treatment with PA in the presence or absence of ETO, the cells were stained with 200µl of 5µM MitoSox^™^ Red following the manufacturer’s protocol. The cells were incubated for 10 mins at 37°C and 5% CO_2_ and then imaged under a fluorescent microscope. Images were captured using Olympus CKX53 inverted fluorescence microscope and using Olympus DP-74 camera.

### 2.6 Quantification of fluorescent intensity

Fluorescent images of MitoSox^™^ stained rTDC were captured from 3 independent experiments. All the images were captured with an Olympus DP-74 camera keeping the exposure time, aperture, and fluorescent intensity constant in Nikon CKX53 microscope. Prior to analysis, images were calibrated, and the mean fluorescent intensity in the red channel was quantified from at least 6 images captured from 3 independent biological experiments.

### 2.7 Assessment of tenocyte mitochondrial respiration

The effects of PA treatment in the presence or absence of ETO on rTDCs mitochondrial function was assessed. rTDCs were grown to confluence and then exposed to one of three treatments: 10 µg/ml PA, 50 µM ETO for 2h before the addition of 10 µg/ml PA, or vehicle control (PBS) for 48h prior to mitochondrial function measurements. Cells were trypsinised, washed in PBS, and resuspended in mitochondrial respiration buffer (MiR05; in mM: 110 sucrose, 60 K-lactobionate, 20 HEPES, 20 taurine, 10 KH_2_PO_4_, 3 MgCl_2_, 0.5 EGTA, and 1 mg/ml fraction V BSA, pH 7.1 at 37°C).

Mitochondrial respiration was measured from 1-2 M tenocytes in an Oroboros O_2_k Oxygraph (Oroboros Instruments, Innsbruck, Austria) in 2 mL MiR05, at 37°C. Routine respiration rate was determined in intact cells, after which cells were permeabilised by the addition of digitonin. Following permeabilisation, respiration was measured following a protocol we have previously described (Hedges et al., 2019). Briefly, malate (0.5 mM) and pyruvate (5 mM) supported complex-I-mediated leak (non-phosphorylating) respiration (CI leak), followed by ADP (2.5 mM) to support oxidative phosphorylation (CI oxphos). Succinate (10 mM) was added to support respiration from complex I and II combined (CI + II oxphos), after which oligomycin (10 nM) was added to measure non-phosphorylation respiration after inhibition of ATP synthase (CI + II leak). Sequential titrations of CCCP (0.5 µM) were added to uncouple electron transport from ATP synthesis and measure the capacity of the electron transport system (ETS). All respiration states were corrected for residual non-mitochondrial oxygen consumption, determined following the addition of antimycin-A (5 µM) to inhibit complex III.

### 2.8 Statistical analysis

GraphPad Prism 8.2.1 (GraphPad Software) was used for statistical analysis. Data from all experiments were analysed using one-way analysis of variance (ANOVA) or two-way ANOVA with post-hoc Tukey’s test or Dunnett’s test, respectively. Tests were 2-tailed, and a 5% significance level was maintained throughout the study.

## 3 Results

### 3.1 Increased concentrations of the saturated fatty acids PA and SA reduce cell viability

rTDCs treated with 10 µg/ml PA had a 15% decrease in cell viability at 24 h, and a 60% reduction in cell viability at 48 h, compared to the control group (Figure 1). Treatment with 10 µg/ml SA decreased the cell viability by 40% at 48 h, compared to the control group. In contrast, OA did not affect rTDC viability. No significant difference in cell viability was observed for all FFA treatments at concentrations below 10 µg/ml.

**Figure 1.**
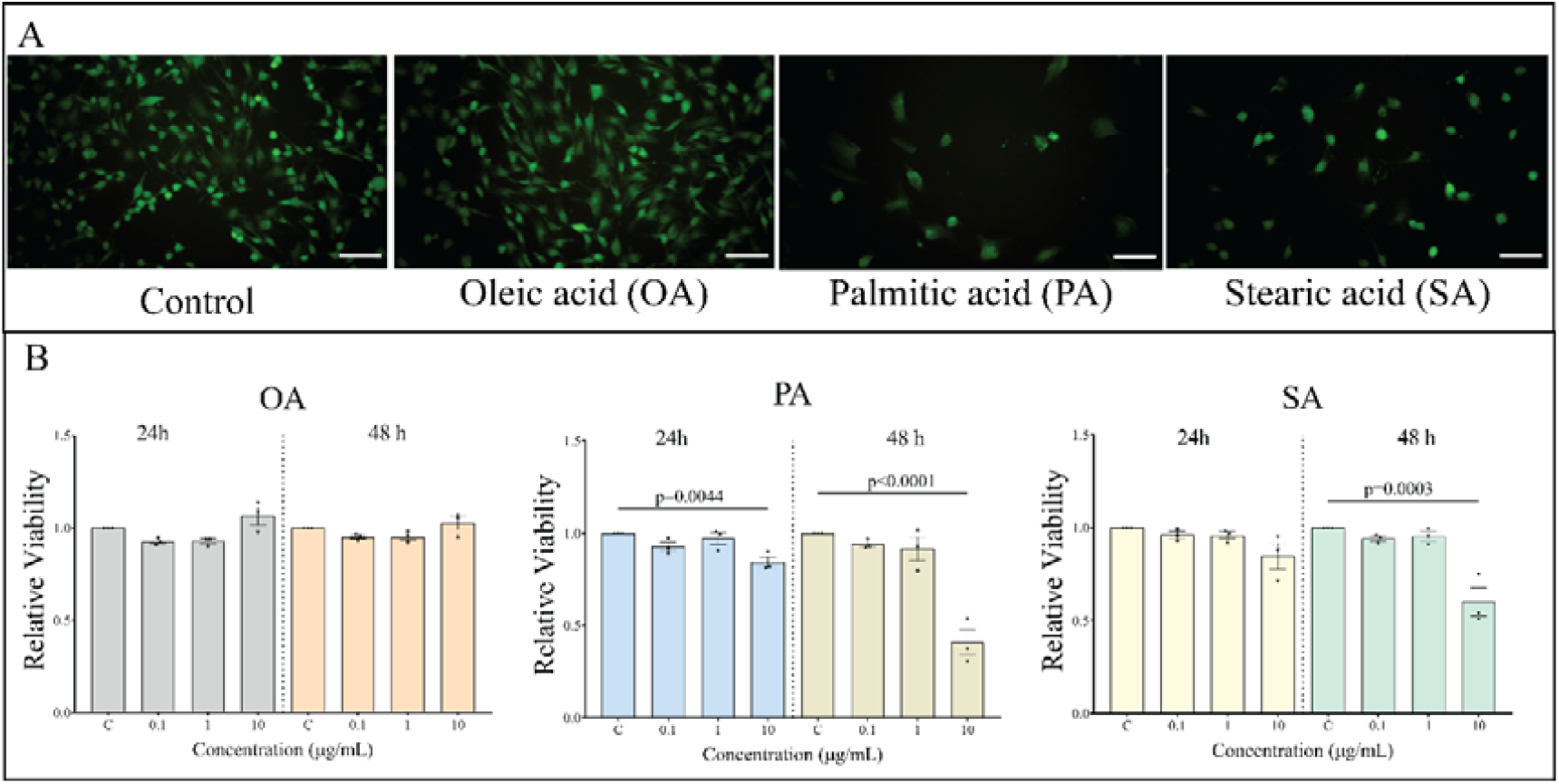
Treatment with saturated FFA decreases rTDC viability A. Representative images of calcein-stained cells after 48h of control, OA, PA, and SA (10µg/ml) treatment. Scale bar100µm. B. Viability of cells treated with OA, PA, and SA for 24 or 48 h. Cell viability was determined by alamarblue™ assay. Results are presented as means ±SEM (n≥3). Groups were compared by one-way ANOVA with post-hoc Dunnett’s test. Comparisons with p values<0.05 are indicated on the graph.

### 3.2 PA inhibits the expression of tenocyte marker genes and induces the expression of genes related to matrix degradation and inflammation

We studied the effects of OA, PA, and SA on the expression of *Scx, Tnmd, Col1a1, Runx2, Sox9, Mmp3, Mmp13*, and *Ptgs2* (COX-2) in rTDCs. The expression of tendon differentiation markers was inhibited by 10 µg/ml of PA: at 24 h, *Scx* expression was approximately 4-fold lower than in the control cells, and at 48 h, the expression of *Tnmd* was approximately 3.5-fold lower than in the control cells (Figure 2). The expression levels of *Mmp3, Mmp13*, and *Ptgs2* were approximately 12, 26, and 9-fold higher than the control, respectively, in cells treated with 10 µg/ml of PA for 48 h. OA had no effect on the expression of the eight genes tested here, and 10 µg/ml of SA only slightly reduced the expression of *Sox9* at 48 h (supplementary Figure 1, 2).

**Figure 2.**
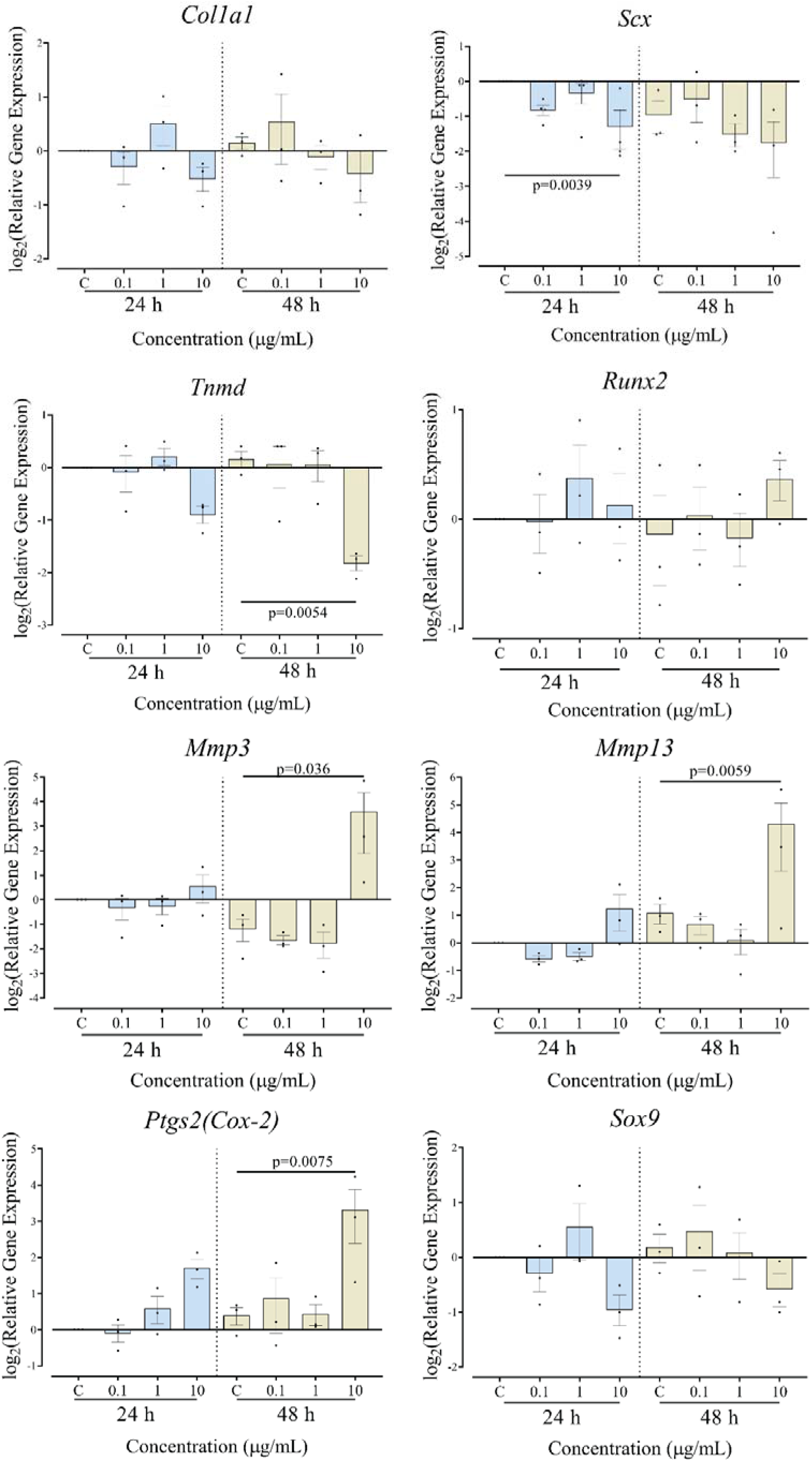
Gene expression of markers of differentiation, inflammation, and matrix degradation in rTDC treated with PA rTDCs were treated with 0.1, 1, and 10 µg/ml PA for 24h and 48h. Expression levels are represented relative to control cells at 24h. Each dot represents one biological experiment, means ± SEM are indicated. Groups were compared by two-way ANOVA with post-hoc Dunnett’s test. Comparisons with p values<0.05 are indicated on the graph.

### 3.3 OA, PA, and SA increase *Cpt1* and *Pdk4* expression in rTDCs

The expression of genes encoding two mitochondrial enzymes: *Cpt1*, encoding the carnitine palmitoyltransferase I (CPT1), and *Pdk4*, encoding pyruvate dehydrogenase kinase 4 (PDK4) increased in rTDCs treated with OA, PA, and SA. In general, in rTDCs treated with 10 µg/ml FFA, the expression of *Cpt1* was 5–10-fold higher than in the controls, whereas the expression of *Pdk4* increased 20–50-fold, and the level of expression was similar at 24 and 48 h (Figure 3). OA, at 10 µg/ml, increased the expression of *Cpt1* and *Pdk4* at the two time points, whereas the effects of PA and SA were somewhat less consistent.

**Figure 3.**
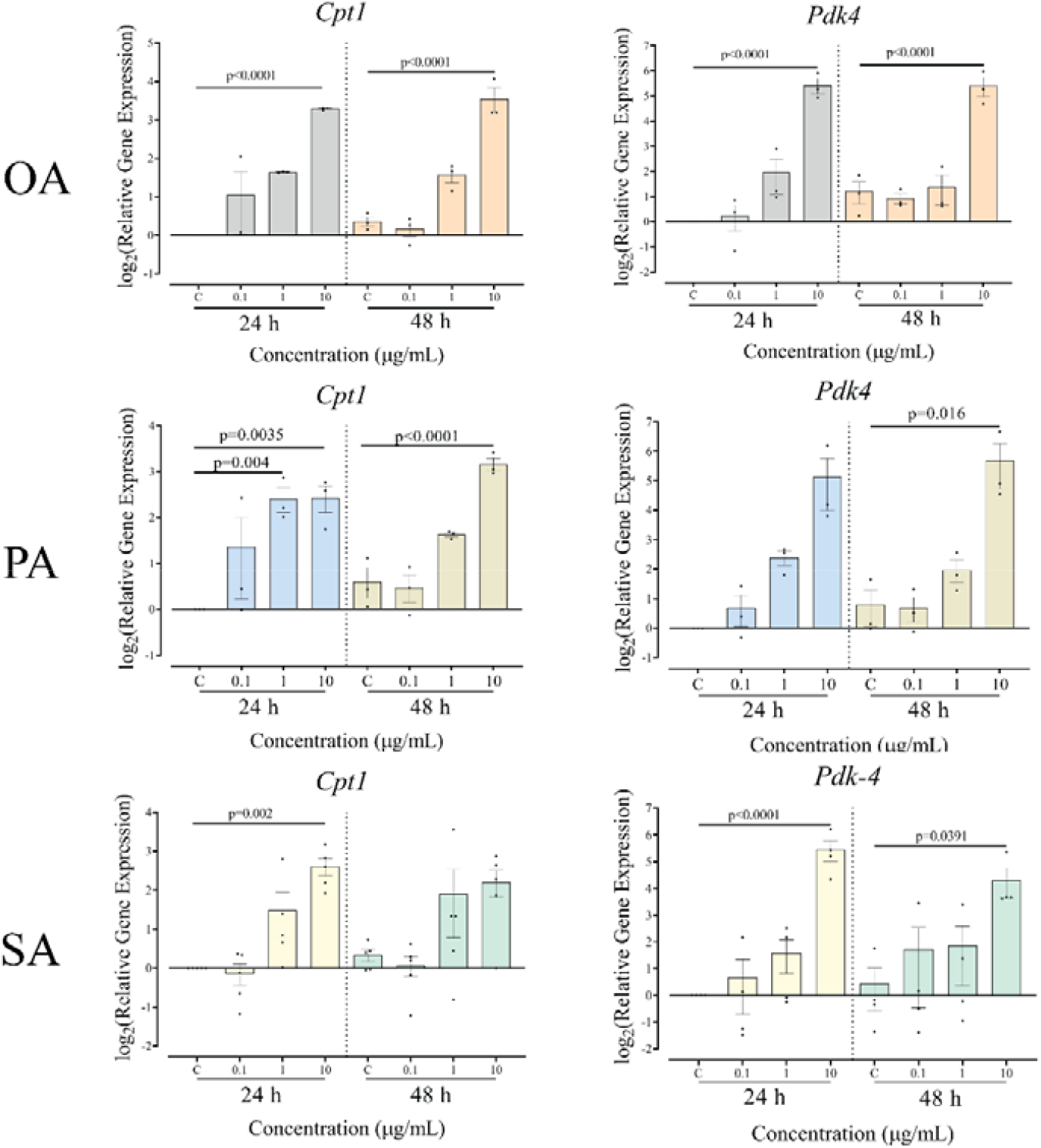
Expression of Cpt1 and Pdk4 genes in rTDCs at 24h and 48h of treatment with OA, PA, and SA. rTDC were treated with 0.1, 1, and 10µg/ml of the fatty acids. Expression levels are represented relative to control cells at 24h. Each dot represents one biological experiment, means ± SEM are indicated (n≥3). Groups were compared by two-way ANOVA with post-hoc Dunnett’s test. Comparisons with p values<0.05 are indicated on the graph.

### 3.4 Etomoxir treatment partially attenuates PA-induced negative changes in rTDC

#### 3.4.1 Etomoxir treatment attenuates the effect of PA on rTDC viability

Treatment with 50 µM ETO for 2 h before the addition of 10 µg/ml PA attenuated the inhibitory effect of PA on rTDC viability (Figure 4). Viability was 53% of the control in rTDCs treated with 10 µg/ml of PA for 48 h, and pre-treatment with ETO attenuated the PA effect by 53%, resulting in cell viability of 72% of the control level.

**Figure 4.**
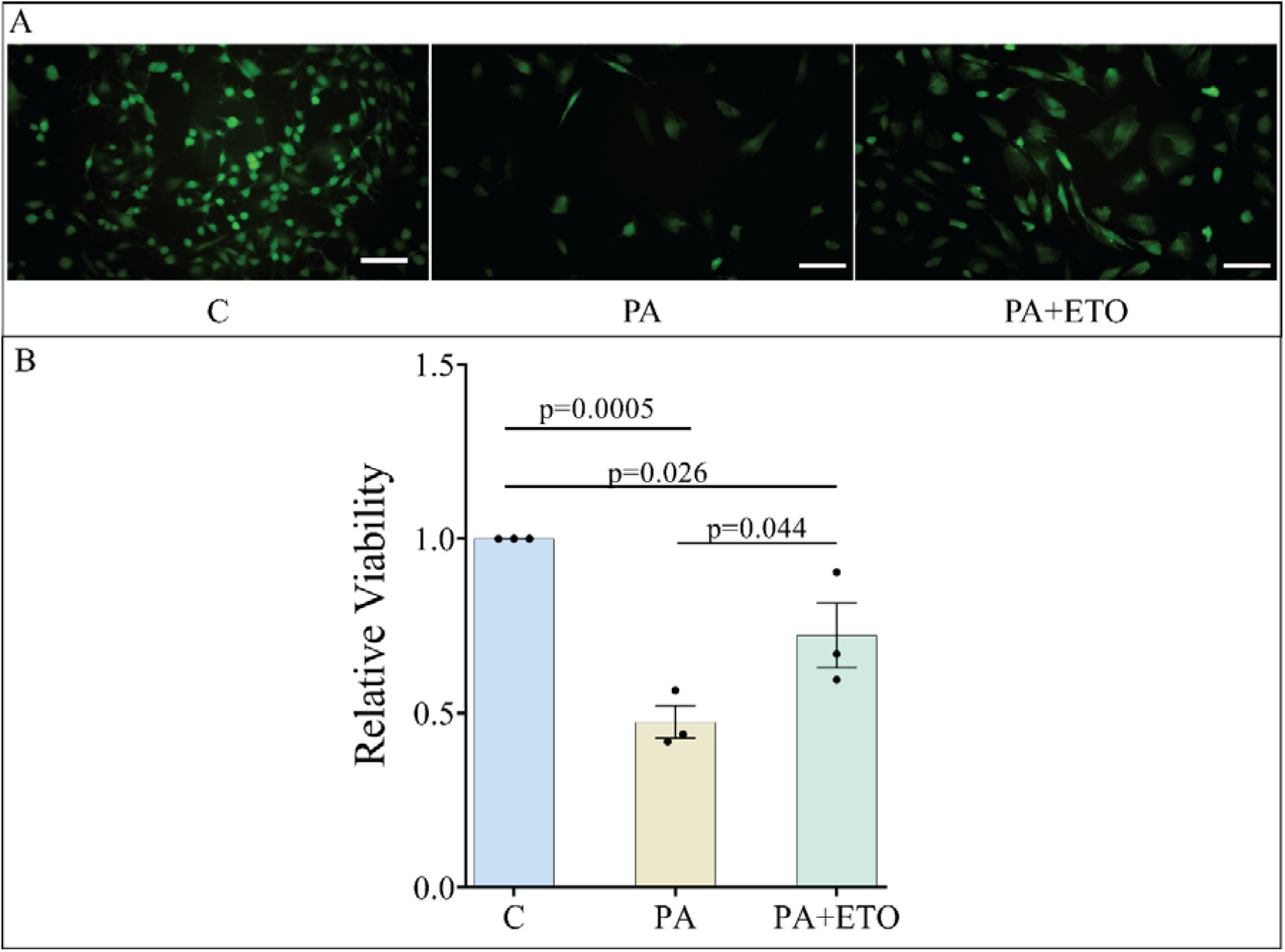
Viability of cells treated with ETO and PA A. Representative image of calcein-stained cells after 48h treatment with control, 10g/ml PA, and PA+ETO. B. Cell viability at 48h was determined by alamarblue™ assay. Results are presented as means±SEM (n=3). Groups were compared by one-way ANOVA with post-hoc Dunnett’s test. C, control cells; PA, 10µg/ml PA treatment; PA+ETO, 50µM etomoxir treatment followed by 10µg/ml PA treatment

#### 3.4.2 Etomoxir treatment partially attenuates the PA-induced effects on rTDC gene expression

Pre-treatment with 50 µg/ml ETO for two hours before the addition of 10 µg/ml PA for 48 h attenuated the PA-induced increase in the expression of *Mmp3* and *Pdk4*. In comparison to control cells, *Mmp3* expression was over 15-fold higher in cells treated with PA, and only 5-fold higher in the presence of ETO; the expression of Pdk4 was reduced from75-fold over the control to 38-fold in the presence of ETO. Pre-treatment with ETO did not change the expression of *Tnmd, Scx, Mmp13, Ptgs2*, and *Cpt1* in rTDCs (Figure 5).

**Figure 5.**
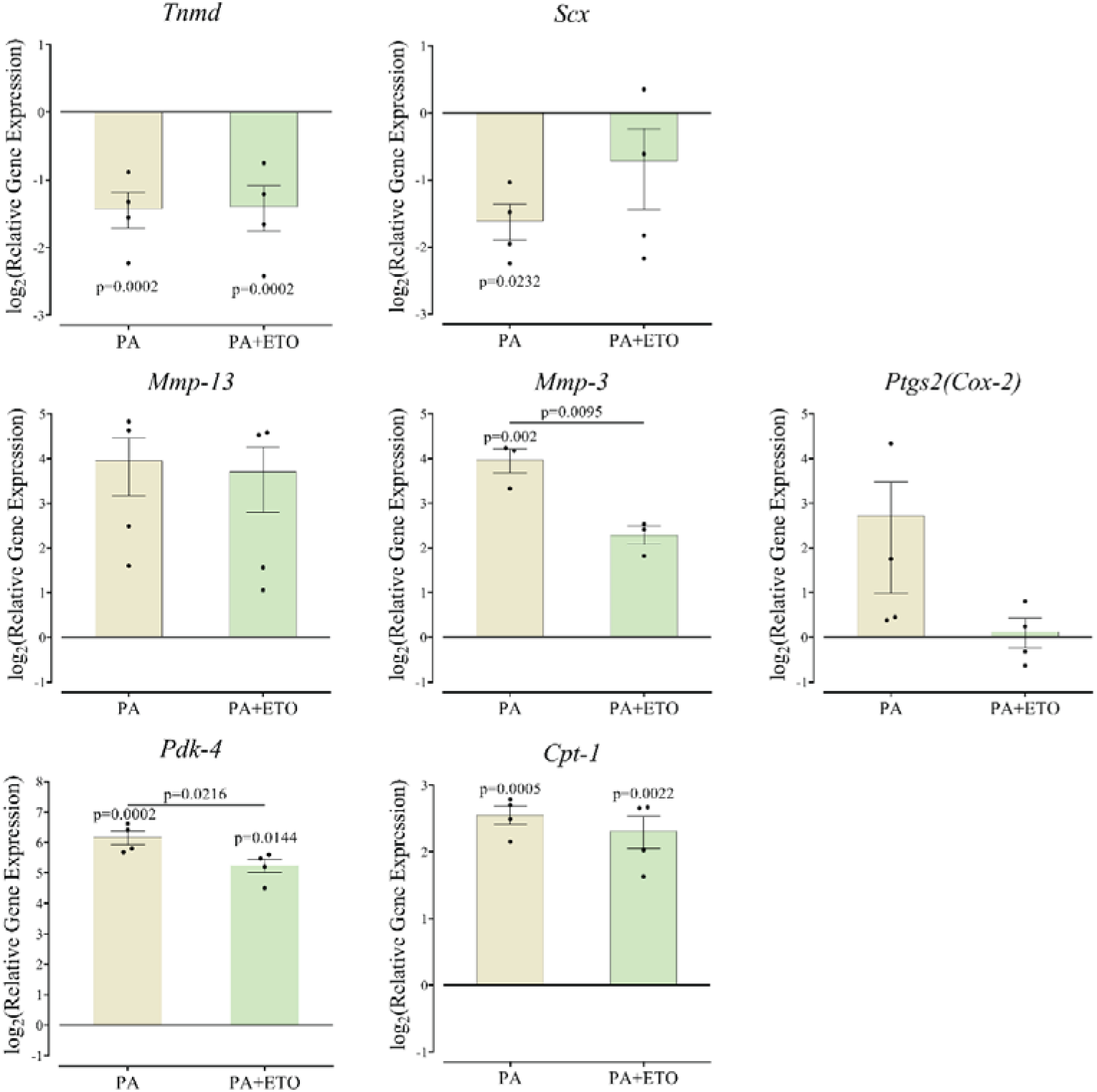
Relative gene expression in rTDC treated with PA or PA+ETO. rTDC were treated with 10µg/ml PA for 48h with or without pre-treatment with 50µM ETO. Expression levels are expressed relative to control cells (line at y=0). Results are represented as means ±SEM (n=4). Groups were compared by one-way ANOVA with post-hoc Tukey’s test. Comparisons with p values<0.05 are indicated on the graph; p values above the individual bars indicate comparisons to control, untreated cells.

### 3.5 PA treatment increased ROS production and respiration

MitoSox staining demonstrated that ROS production increased in rTDCs treated with 10µg/ml PA for 48 h, with the mean fluorescent intensity approximately 40% higher than the control group (Fig. 6A, B). Pre-treatment with ETO blocked the PA-induced increase in ROS production.

**Figure 6.**
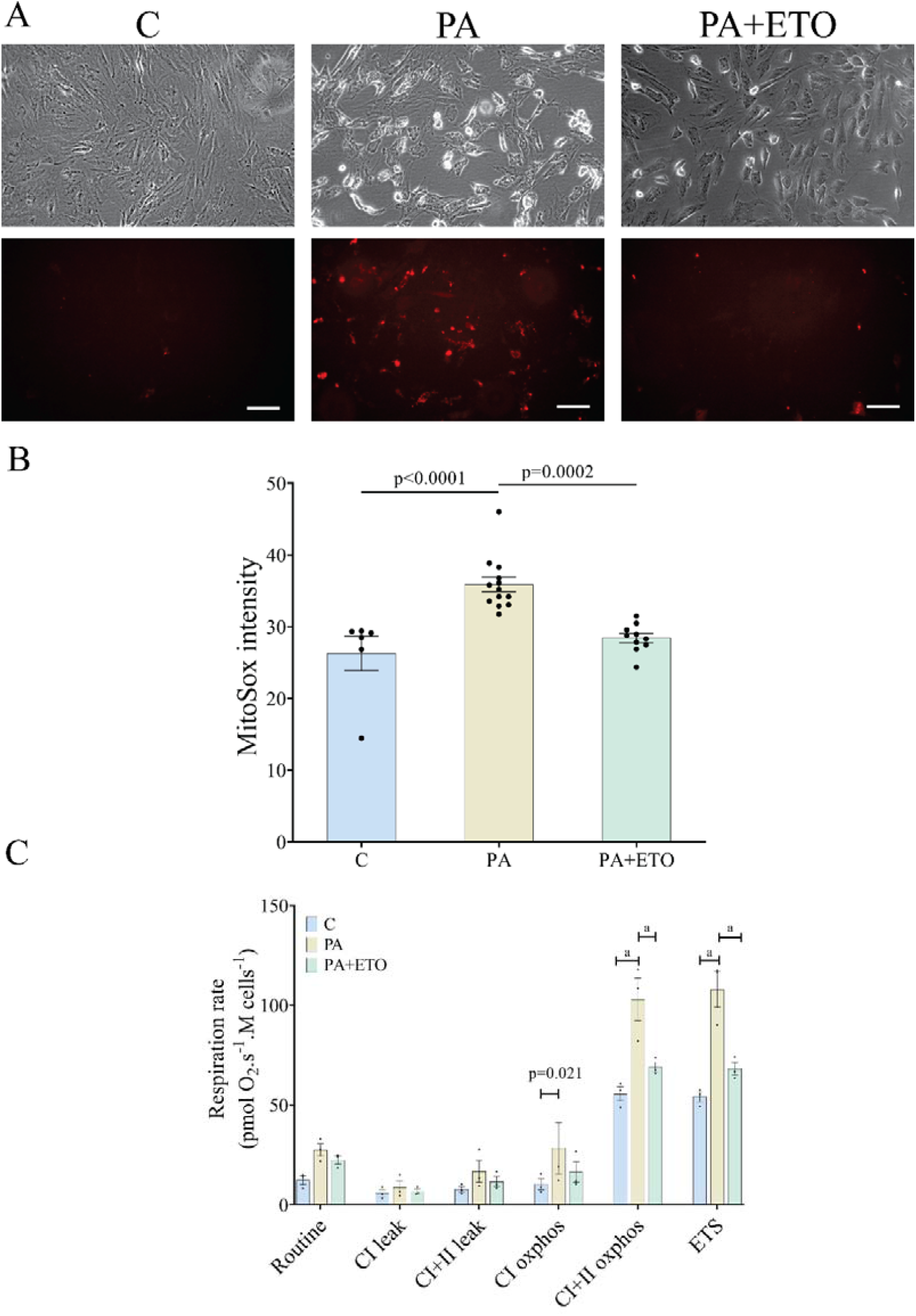
Treatment of rTDCs with PA increased ROS generation and mitochondrial respiration. A. Representative images of rTDC in brightfield and stained with MitoSox™ red. Treatment with PA for 48h increased ROS production by the rTDC. (Scale bar=100µm). B. Quantification of MitoSox™ red Fluorescent intensity. Dots represent individual data points, means ± SEM are presented. Data were compared by one-way ANOVA with post-hoc Dunnett’s test. C. rTDC mitochondrial function when treated with PA or PA+ETO under different conditions. Dots represent individual data points, means ± SEM are presented. Data were compared by two-way ANOVA with post-hoc Fisher’s LSD test. ‘a’ represents p<0.0001 between the bars.

Mitochondrial oxygen consumption was almost 2-fold higher during CI+II oxphos and ETS states in rTDCs treated with 10µg/ml PA for 48h compared to control cells (Fig. 6C). Similar to ROS production, pre-treatment with 50µM ETO blocked the PA-induced increase in mitochondrial respiration under CI+II oxphos and ETS.

## 4 Discussion

We found that the two saturated FFA, PA and SA, reduced rTDC viability. PA, but not SA, inhibited the expression of tendon marker genes and induced the expression of inflammatory markers. The unsaturated FFA, OA, did not affect cell viability and the expression of these genes. The expression of *Cpt1* and *Pdk4* were greatly induced by all three FFA, indicating an increase in mitochondrial fatty acid oxidation. PA was also found to increase ROS generation and respiration in rTDCs. Blocking the transport of PA into the mitochondria attenuated its effects on cell viability, gene expression, ROS generation, and respiration, suggesting that PA effects in rTDC are partially mediated by mitochondrial processing.

We have previously shown that diet-induced obesity in rats impairs rotator cuff tendon healing and alters tendon cell phenotype (Bolam et al., 2021; Bolam et al., 2022). However, the specific contribution of obesity-associated increases in circulating glucose, cholesterol, and FFA to the tendon phenotype and function has not been analysed in these studies. In general, investigations of the mechanisms that associate obesity and tendinopathy focused mostly on the effects of high cholesterol and glucose levels on tendons (Longo et al., 2009; Tilley et al., 2015). High glucose concentrations increased rTDC apoptosis in vitro and altered the microstructure of rat Achilles tendons in vivo (Poulsen et al., 2011; Ueda et al., 2018; Wu et al., 2017). Clinical studies have found that increased cholesterol levels are a risk factor for tendon injury (Mathiak et al., 1999; Ozgurtas et al., 2003; Tilley et al., 2015). High cholesterol levels were shown to inhibit tendon marker expression and increase ROS production (Gunaratnam et al., 2013). The contribution of the obesity-related increase in circulating FFA to tendinopathy has not been investigated previously.

The concentrations of FFA in tendon tissue are unknown. In the circulation, the concentrations of FFA are typically about 400µg/ml in non-obese people and approximately 1.5-fold higher in people with obesity (Kim et al., 2010; Ni et al., 2015; Abdelmagid et al., 2015). FFA concentrations in tendons are expected to be much lower because of the avascular nature of tendon tissue and the limited transport of nutrients from the circulation (Tempfer and Traweger, 2015). In the current study, we tested PA, SA, and OA at concentrations of 0.1, 1, and 10 µg/ml, and found that at 10 µg/ml, PA had negative effects on cell viability and gene expression. Previous studies found that increased PA concentrations caused the death of skeletal muscle cells and pancreatic ß-cells(Ahn et al., 2013; Feng et al., 2014; Gunaratnam et al., 2013; Nemecz et al., 2019). Our finding that OA does not affect cell viability is also consistent with previous studies that have shown OA did not increase cell death and even had a protective effect against PA treatment(Ahn et al., 2013). We hypothesise that the obesity-associated increase in circulating PA elevates the local concentrations of PA in tendon tissue to a level that is detrimental to tendon cells, as observed in our in vitro model.

Treatment with 10 µg/ml PA modified gene expression in rTDCs. *Scx* and *Tnmd* are the two main tendon marker genes, and their reduced expression in PA-treated rTDCs indicates the dedifferentiation of these specialised cells. We also found that treatment of rTDCs with 10 µg/ml PA induced the expression of *Mmp3, Mmp13*, and *Ptgs2*. Previous studies have shown that increased FFA concentration induce the expression of MMPs and the secretion of PGE-2 (prostaglandin E-2) from immune cells (Hellmann et al., 2013; Hu et al., 2016; Sindhu et al., 2016). The increased expression of *Ptgs2* in PA-treated rTDCs suggests the activation of inflammatory pathways mediated by elevated COX-2 and the secretion of the end product of the COX-2 pathway, PGE-2. MMP enzymes degrade extracellular matrix and play an important role in tissue homeostasis and remodelling. MMP-1, 3, and 13 contribute to the degradation of type-I and type-III collagen, the two main collagen types found in tendon extracellular matrix (Riley et al., 1994). Increased MMP expression was observed in tendinopathy(De Mos et al., 2007; Riley, 2004) and in rTDCs isolated from diet-induced obese rats (Bolam et al., 2022). The increase in MMP-13 expression can indicate the degradation of the tendon matrix and deterioration of tendon biomechanical properties.

To understand if FFA treatment alters metabolic pathways in rTDCs, we investigated the effect of FFA treatment on two primary markers of the fatty acid metabolism pathway: *Cpt1* and *Pdk4*. CPT-1 regulates fatty acid transport into the mitochondria (Slawik and Vidal-Puig, 2006), and PDK-4 inhibits pyruvate dehydrogenase, therefore impairing glycolysis (Pettersen et al., 2019). Increased *Cpt1* expression upon FFA treatment, which was consistent for the three FFAs used in this study, could potentially indicate increased lipid metabolism. It is important to note that OA increased the expression of *Cpt1* and *Pdk4* but did not affect cell viability or the expression of any of the other genes tested, indicating that activation of this pathway is insufficient to produce the other effects in TDCs.

Previous studies have shown that increased FFA metabolism is associated with increased ROS production in the mitochondria (Schönfeld and Wojtczak, 2007; Su et al., 2019). High concentrations of ROS cause oxidative damage and lead to cell death (Simon et al., 2000). Here, we found that PA increased ROS production in rTDCs. To determine whether mitochondrial processing mediates the effects of PA in rTDC, we used ETO to block PA transport into the mitochondria. ETO is a competitive inhibitor that binds CPT-1 with a higher affinity than its physiological substrate, palmitoyl CoA (Schlaepfer et al., 2014; Xu et al., 2003). We found that ETO blocked the PA-mediated increase in ROS production but only attenuated the negative effect of PA on rTDC viability, indicating that additional pathways are involved in mediating this effect. We also observed an increase in mitochondrial respiration, which, in conjunction with increased ROS production, suggests that PA treatment increases mitochondrial content, rather than altering intrinsic properties of rTDC mitochondria. Our finding that respiration was increased during CI+II Oxphos is consistent with previous data showing that knockdown of *Scx* resulted in increased expression of genes relating to oxidative phosphorylation (Nichols et al., 2018). Thus, the PA-induced increase in mitochondrial respiration seen here may be secondary to a decrease in *Scx* expression.

Our study had several limitations. The concentrations of FFA we used were based on estimates, because the concentrations of FFA in tendon tissue are currently unknown. We aimed to model the avascular tendon environment by using concentrations that were lower than those reported for human circulation. When knowledge of the local tendon FFA concentrations becomes available, it would allow us to adjust the model and accurately reproduce the conditions in tendons of non-obese people and people with obesity. Another limitation of the study is that the effects of FFA on gene expression in rTDCs were not corroborated by analysis of the corresponding protein levels. However, the investigations of cell viability, ROS production, and respiration provide complementary functional evidence for the effects of FFAs in rTDCs. In future studies, it would be interesting to investigate the mechanisms that underlie the differential effects of OA, SA, and PA identified here.

Our study found that PA reduces cell viability, alters gene expression, and increases ROS production and respiration in rTDCs. The changes in gene expression are consistent with the typical changes observed in tendinopathy, including cellular dedifferentiation, increased extracellular matrix degradation, and the development of inflammatory conditions. Mitochondrial processing is involved in mediating the effects of PA in rTDCs. Our study suggests that the high concentrations of circulating FFA is one of the mediators of the increased risk for tendinopathy in people with obesity.

## 5 Acknowledgement

This work was supported by the Auckland Medical Research Foundation #1117017 project grant and senior research fellowship (DM).

## Reference

Abate, M., Schiavone, C., Di Carlo, L. and Salini, V. (2012). Achilles tendon and plantar fascia in recently diagnosed type II diabetes: Role of body mass index. Clin. Rheumatol. 31, 1109–1113.

Abdelmagid, S. A., Clarke, S. E., Nielsen, D. E., Badawi, A., El-Sohemy, A., Mutch, D. M. and Ma, D. W. L. (2015). Comprehensive Profiling of Plasma Fatty Acid Concentrations in Young Healthy Canadian Adults. PLoS One 10, e0116195.

Ahn, J. H., Kim, M. H., Kwon, H. J., Choi, S. Y. and Kwon, H. Y. (2013). Protective Effects of Oleic Acid Against Palmitic Acid-Induced Apoptosis in Pancreatic AR42J Cells and Its Mechanisms. Korean J. Physiol. Pharmacol. 17, 43.

Andarawis-Puri, N., Flatow, E. L. and Soslowsky, L. J. (2015). Tendon basic science: Development, repair, regeneration, and healing. J. Orthop. Res. 33, 780–784.

Bi, X., Yeo, P. L. Q., Loo, Y. T. and Henry, C. J. (2019). Associations between circulating fatty acid levels and metabolic risk factors. J. Nutr. Intermed. Metab. 15, 65–69.

Boden, G. (2008). Obesity and Free Fatty Acids. Endocrinol. Metab. Clin. North Am. 37, 635–646.

Bolam, S. M., Park, Y. E., Konar, S., Callon, K. E., Workman, J., Monk, A. P., Coleman, B., Cornish, J., Vickers, M. H., Munro, J. T., et al. (2021). Obesity Impairs Enthesis Healing After Rotator Cuff Repair in a Rat Model. Am. J. Sports Med. 49, 3959–3969.

Bolam, S. M., Konar, S., Park, Y.-E., Callon, K. E., Workman, J., Monk, A. P., Coleman, B., Cornish, J., Vickers, M. H., Munro, J. T., et al. (2022). A high-fat diet has negative effects on tendon resident cells in an in vivo rat model. Int. Orthop. 46, 1181–1190.

Chhana, A., Callon, K. E., Dray, M., Pool, B., Naot, D., Gamble, G. D., Coleman, B., McCarthy, G., McQueen, F. M., Cornish, J., et al. (2014). Interactions between tenocytes and monosodium urate monohydrate crystals: Implications for tendon involvement in gout. Ann. Rheum. Dis. 73, 1737–1741.

Cornish, J., MacGibbon, A., Lin, J.-M., Watson, M., Callon, K. E., Tong, P. C., Dunford, J. E., van der Does, Y., Williams, G. A., Grey, A. B., et al. (2008). Modulation of osteoclastogenesis by fatty acids. Endocrinology 149, 5688–95.

De Mos, M., Van El, B., Degroot, J., Jahr, H., Van Schie, H. T. M., Van Arkel, E. R., Tol, H., Heijboer, R., Van Osch, G. J. V. M. and Verhaar, J. A. N. (2007). Achilles tendinosis: Changes in biochemical composition and collagen turnover rate. Am. J. Sports Med. 35, 1549–1556.

Elbaz, A., Wu, X., Rivas, D., Gimble, J. M. and Duque, G. (2010). Inhibition of fatty acid biosynthesis prevents adipocyte lipotoxicity on human osteoblasts in vitro. J. Cell. Mol. Med. 14, 982–991.

Feng, X., Wang, T., Leng, J., Chen, Y., Liu, J., Liu, Y., Wang, J. and Phosphorylation, S. (2014). Palmitate Contributes to Insulin Resistance through Downregulation of the Src-Mediated Phosphorylation of Akt in C2C12 Myotubes Palmitate Contributes to Insulin Resistance through Downregulation. 8451,.

Gunaratnam, K., Vidal, C., Boadle, R., Thekkedam, C. and Duque, G. (2013). Mechanisms of palmitate-induced cell death in human osteoblasts. Biol. Open 2, 1382–1389.

Hedges, C. P., Woodhead, J. S. T., Wang, H. W., Mitchell, C. J., Cameron-Smith, D., Hickey, A. J. R. and Merry, T. L. (2019). Peripheral blood mononuclear cells do not reflect skeletal muscle mitochondrial function or adaptation to high-intensity interval training in healthy young men. J. Appl. Physiol. 126, 454–461.

Hellmann, J., Zhang, M. J., Tang, Y., Rane, M., Bhatnagar, A. and Spite, M. (2013). Increased Saturated Fatty Acids in Obesity Alter Resolution of Inflammation in Part by Stimulating Prostaglandin Production. J. Immunol. 191, 1383–1392.

Hu, X., Cifarelli, V., Sun, S., Kuda, O., Abumrad, N. A. and Su, X. (2016). Major role of adipocyte prostaglandin E2 in lipolysis-induced macrophage recruitment. J. Lipid Res. 57, 663.

Kim, J. Y., Park, J. Y., Kim, O. Y., Ham, B. M., Kim, H. J., Kwon, D. Y., Jang, Y. and Lee, J. H. (2010). Metabolic profiling of plasma in overweight/obese and lean men using ultra performance liquid chromatography and Q-TOF mass spectrometry (UPLC-Q-TOF MS). J. Proteome Res. 9, 4368–4375.

Li, K., Deng, Y., Deng, G., Chen, P., Wang, Y., Wu, H., Ji, Z., Yao, Z., Zhang, X., Yu, B., et al. (2020). High cholesterol induces apoptosis and autophagy through the ROS-activated AKT/FOXO1 pathway in tendon-derived stem cells. Stem Cell Res. Ther. 11, 1–16.

Lin, Y., Li, Y., Rui, Y., Dai, G., Shi, L., Xu, H.-L., Ni, M., Zhao, S., Chen, H., Wang, C., et al. (2017). The effects of high glucose on tendon-derived stem cells: implications of the pathogenesis of diabetic tendon disorders. Oncotarget 8, 17518–17528.

Listenberger, L. L., Han, X., Lewis, S. E., Cases, S., Farese, R. V., Ory, D. S. and Schaffer, J. E. (2003). Triglyceride accumulation protects against fatty acid-induced lipotoxicity. Proc. Natl. Acad. Sci. U. S. A. 100, 3077–3082.

Longo, U. G., Franceschi, F., Ruzzini, L., Spiezia, F., Maffulli, N. and Denaro, V. (2009). Higher fasting plasma glucose levels within the normoglycaemic range and rotator cuff tears. Br. J. Sports Med. 43, 284–7.

Mathiak, G., Wening, J. V., Mathiak, M., Neville, L. F. and Jungbluth, K. H. (1999). Serum cholesterol is elevated in patients with Achilles tendon ruptures. Arch. Orthop. Trauma Surg. 119, 280–284.

Musson, D. S., Naot, D., Chhana, A., Callon, K. E., Choi, A. J., Matthews, B. G., Dunbar, P. R., Cornish, J., Coleman, B., McIntosh, J. D., et al. (2015). In Vitro Evaluation of a Novel Non-Mulberry Silk Scaffold for Use in Tendon Regeneration. Tissue Eng. Part A 21, 1539–1551.

Nemecz, M., Constantin, A., Dumitrescu, M., Alexandru, N., Filippi, A., Tanko, G. and Georgescu, A. (2019). The distinct effects of palmitic and oleic acid on pancreatic beta cell function: The elucidation of associated mechanisms and effector molecules. Front. Pharmacol. 9, 1–16.

Ni, Y., Zhao, L., Yu, H., Ma, X., Bao, Y., Rajani, C., Loo, L. W. M., Shvetsov, Y. B., Yu, H., Chen, T., et al. (2015). Circulating Unsaturated Fatty Acids Delineate the Metabolic Status of Obese Individuals. EBioMedicine 2, 1513–1522.

Nichols, A. E. C., Settlage, R. E., Werre, S. R. and Dahlgren, L. A. (2018). Novel roles for scleraxis in regulating adult tenocyte function. BMC Cell Biol. 19,.

Ozgurtas, T., Yildiz, C., Serdar, M., Atesalp, S. and Kutluay, T. (2003). Is high concentration of serum lipids a risk factor for Achilles tendon rupture? Clin. Chim. Acta 331, 25–28.

Pettersen, I. K. N., Tusubira, D., Ashrafi, H., Dyrstad, S. E., Hansen, L., Liu, X. Z., Nilsson, L. I. H., Løvsletten, N. G., Berge, K., Wergedahl, H., et al. (2019). Upregulated PDK4 expression is a sensitive marker of increased fatty acid oxidation. Mitochondrion 49, 97–110.

Poulsen, R. C., Carr, A. J. and Hulley, P. A. (2011). Protection against Glucocorticoid-Induced Damage in Human Tenocytes by Modulation of ERK, Akt, and Forkhead Signaling. Endocrinology 152, 503–514.

Poulsen, R. C., Knowles, H. J., Carr, A. J. and Hulley, P. A. (2014). Cell differentiation versus cell death: extracellular glucose is a key determinant of cell fate following oxidative stress exposure. Cell Death Dis. 5, e1074–e1074.

Reaven, G. M. (1995). Pathophysiology of insulin resistance in human disease. Physiol. Rev. 75, 473–486.

Rechardt, M., Shiri, R., Karppinen, J., Jula, A., Heliövaara, M. and Viikari-Juntura, E. (2010). Lifestyle and metabolic factors in relation to shoulder pain and rotator cuff tendinitis: A population-based study. BMC Musculoskelet. Disord. 11, 165.

Riley, G. (2004). The pathogenesis of tendinopathy. A molecular perspective. Rheumatology 43, 131–142.

Riley, G. P., Harrall, H., Constant, C. R., Chard, M. D., Cawston, T. E. and Hazleman, B. L. (1994). Tendon degeneration and chronic shoulder pain: Changes in the collagen composition of the human rotator cuff tendons in rotator cuff tendinitis. Ann. Rheum. Dis.

Rosca, M. G., Vazquez, E. J., Chen, Q., Kerner, J., Kern, T. S. and Hoppel, C. L. (2012). Oxidation of Fatty Acids Is the Source of Increased Mitochondrial Reactive Oxygen Species Production in Kidney Cortical Tubules in Early Diabetes. Diabetes 61, 2074.

Schlaepfer, I. R., Rider, L., Rodrigues, L. U., Gijón, M. A., Pac, C. T., Romero, L., Cimic, A., Sirintrapun, S. J., Glodé, L. M., Eckel, R. H., et al. (2014). Lipid catabolism via CPT1 as a therapeutic target for prostate cancer. Mol. Cancer Ther. 13, 2361–2371.

Schönfeld, P. and Wojtczak, L. (2007). Fatty acids decrease mitochondrial generation of reactive oxygen species at the reverse electron transport but increase it at the forward transport. Biochim. Biophys. Acta - Bioenerg. 1767, 1032–1040.

Simon, H. U., Haj-Yehia, A. and Levi-Schaffer, F. (2000). Role of reactive oxygen species (ROS) in apoptosis induction. Apoptosis 2000 55 5, 415–418.

Sindhu, S., Al-Roub, A., Koshy, M., Thomas, R. and Ahmad, R. (2016). Palmitate-Induced MMP-9 Expression in the Human Monocytic Cells is Mediated through the TLR4-MyD88 Dependent Mechanism. Cell. Physiol. Biochem. 39, 889–900.

Slawik, M. and Vidal-Puig, A. J. (2006). Lipotoxicity, overnutrition and energy metabolism in aging. Ageing Res. Rev.

Snezhkina, A. V., Kudryavtseva, A. V., Kardymon, O. L., Savvateeva, M. V., Melnikova, N. V., Krasnov, G. S. and Dmitriev, A. A. (2020). ROS generation and antioxidant defense systems in normal and malignant cells. Oxid. Med. Cell. Longev. 2019,.

Su, L. J., Zhang, J. H., Gomez, H., Murugan, R., Hong, X., Xu, D., Jiang, F. and Peng, Z. Y. (2019). Reactive Oxygen Species-Induced Lipid Peroxidation in Apoptosis, Autophagy, and Ferroptosis. Oxid. Med. Cell. Longev. 2019,.

Taş, S., Yilmaz, S., Onur, M. R., Soylu, A. R., Altuntaş, O. and Korkusuz, F. (2017). Patellar tendon mechanical properties change with gender, body mass index and quadriceps femoris muscle strength. Acta Orthop. Traumatol. Turc. 51, 54–59.

Tempfer, H. and Traweger, A. (2015). Tendon vasculature in health and disease. Front. Physiol. 6, 1–7.

Tilley, B. J., Cook, J. L., Docking, S. I. and Gaida, J. E. (2015). Is higher serum cholesterol associated with altered tendon structure or tendon pain? A systematic review. Br. J. Sports Med. 49, 1504.

Titchener, A. G., Fakis, A., Tambe, A. A., Smith, C., Hubbard, R. B. and Clark, D. I. (2013). Risk factors in lateral epicondylitis (tennis elbow): A case-control study. J. Hand Surg. Eur. Vol. 38, 159–164.

Ueda, Y., Inui, A., Mifune, Y., Sakata, R., Muto, T., Harada, Y., Takase, F., Kataoka, T., Kokubu, T. and Kuroda, R. (2018). The effects of high glucose condition on rat tenocytes in vitro and rat Achilles tendon in vivo. Bone Jt. Res. 7, 362–372.

Wu, Y. F., Wang, H. K., Chang, H. W., Sun, J., Sun, J. S. and Chao, Y. H. (2017). High glucose alters tendon homeostasis through downregulation of the AMPK/Egr1 pathway. Sci. Reports 2017 71 7, 1–12.

Wu, Y. F., Huang, Y. T., Wang, H. K., Yao, C. C. J., Sun, J. S. and Chao, Y. H. (2018). Hyperglycemia Augments the Adipogenic Transdifferentiation Potential of Tenocytes and Is Alleviated by Cyclic Mechanical Stretch. Int. J. Mol. Sci. 19, 90.

Xu, F. Y., Taylor, W. A., Hurd, J. A. and Hatch, G. M. (2003). Etomoxir mediates differential metabolic channeling of fatty acid and glycerol precursors into cardiolipin in H9c2 cells. J. Lipid Res. 44, 415–423.

